# Mechanistic macroecology: exploring the drivers of latitudinal variation in terrestrial body size in a General Ecosystem Model

**DOI:** 10.1101/775957

**Authors:** Michael Brian James Harfoot, Andrew Abraham, Derek P Tittensor, Gabriel C Costa, Søren Faurby, Anat Feldman, Yuval Itescu, Shai Meiri, Ignacio Morales-Castilla, Brunno F Oliveira, Drew Purves

## Abstract

Many mechanisms have been hypothesized to explain Bergmann’s rule - the correlation of body size with latitude. However, it is not feasible to assess the contribution of hypothesised mechanisms by experimental manipulation or statistical correlation. Here, we evaluate two of the principal hypothesised mechanisms, related to thermoregulation and resource availability, using structured experiments in a mechanistic global ecosystem model. We simulated the broad structure of assemblages and ecosystems using the Madingley model, a mechanistic General Ecosystem Model (GEM). We compared emergent modelled biogeographic patterns in body mass to empirical patterns for mammals and birds. We then explored the relative contribution of thermoregulation and resource availability to body mass clines by manipulating the model’s environmental gradients. Madingley produces body size gradients that are in broad agreement with empirical estimates. Thermoregulation and resource availability were both important controls on body mass for endotherms, but only temperature for ectotherms. Our results suggest that seasonality explains animal body mass patterns through a complex set of mechanisms. Process-based GEMs generate broadly realistic biogeographic body mass patterns. Ecologists can use them in novel ways: to explore causality, or for generating and testing hypotheses for large-scale, emergent ecological patterns. At the same time, macroecological patterns are useful for evaluating mechanistic models. Iteratively developing GEMs, and evaluating them against macroecological patterns, could generate new insights into the complex causes of such patterns.

## INTRODUCTION

### The role of mechanistic models in macroecology

The fields of macroecology and biogeography have traditionally adopted a strong observational and descriptive approach to understanding patterns of species distributions, community composition and biodiversity across space and time (Blackburn et al., 1999). Correlative, statistical approaches have been the primary tools for exploring such data but they are unable to establish causal relationships and can be difficult to transfer across geographic space, time and environmental space (Cabral et al., 2017). Mechanistic ecological models are a new approach that can complement such studies. They represent both the composition of ecosystems and the causal relationships determining that composition. So, they permit targeted exploration of causality in macroecology (Connolly et al., 2017), especially at the large scales at which experimental manipulation is not possible.

### Latitudinal variation in body size and the hypothesised mechanisms

Bergmann’s rule, representing one of the oldest recognized macroecological pattern (Bergmann, 1847), hypothesises that body size is positively correlated with latitude and elevation, with heat conservation increased in larger-bodied animals through a smaller surface-area-to-volume ratio (James 1970, Blackburn et al., 1999). Bergmann’s rule can be considered an empirical generalisation (Mayr, 1956, Meiri, 2011) and has been applied in both inter- and intra-specific studies to a broad range of taxa, including ectotherms, in multiple geographic regions. Amongst endotherms, the majority of mammal and bird species appear to exhibit Bergmann clines (Ashton, 2002; Cardillo, 2002; Freckleton et al., 2003; Meiri et al., 2004; Rodriguez et al., 2006; Ramirez et al., 2008; Rodríguez et al., 2008; Olson et al., 2009; Morales-Castilla et al. 2012a,b; Torres-Romero et al. 2016), though converse clines exist, for example in a group of subterranean rodents (Medina et al., 2007). In contrast, the evidence for ectotherms is highly ambiguous, with some taxa conforming well (Olalla-Tárraga and Rodríguez, 2007), whilst others show inverse clines (Mosseau, 1997; Ashton and Feldmann, 2003; Olalla-Tárraga and Rodríguez, 2007; Adams and Church, 2008; Cvetkovic et al., 2009) and some show no clear pattern (Olalla-Tárraga et al., 2006; Pincheria-Donoso and Meiri, 2013; Feldman and Meiri, 2014, Slavenko et al. 2019). Accordingly, there is debate about whether Bergmann’s rule should be considered for ectotherms at all (Watt et al., 2010; Olalla-Tárraga, 2010). Therefore, we focus on latitudinal variation in the body size of endotherms in this study. Even with this choice and despite over 170 years of scientific research, both the generality and the underlying mechanism(s) of this ecogeographic rule thus remain disputed (James, 1970; Blackburn et al., 1999; Meiri and Dayan, 2003; Pincheria-Donoso, 2010; Meiri, 2011).

Multiple mechanisms have been proposed to explain the observed patterns in body mass (Blackburn et al., 1999; Olalla-Tárraga, 2011) (Table 1). Here we describe two leading mechanisms hypothesised to determine body size clines, which will be the focus of this study.

**Table 1.** Hypothesised mechanisms contributing to the latitudinal gradients in animal body size.

| **Mechanism** | **Description** |
| --- | --- |
| *Chance* | Random ancestral colonisation and subsequent diversification may have led to larger body masses at higher latitudes (Blackburn et al., 1999). Alternatively, selective advantages of traits coupled with body mass may cause the pattern. |
| *Migration ability* | Smaller species are associated with lower dispersal abilities. The different dispersal velocities of larger and smaller bodied species during the retreat of the late Pleistocene ice sheet therefore results in underrepresentation of smaller species in areas from which ice has retreated (Blackburn and Gaston, 1996; Olalla-Tárraga and Rodríguez, 2007; Morales-Castilla et al., 2012a,b; Martinez et al., 2013). |
| *Predation* | Predation risk appears to decrease with distance from the equator. This allows species at high latitude to mature to a larger size (Ashton et al., 2000). |
| *Thermoregulation* | If animals are the same shape, larger species have increased heat conservation in cooler environments thereby augmenting survival and energy allocation to growth (Bergmann, 1847). |
| *Resource availability* | Body mass must be maintained by a sufficient supply of food resources (Rosenzweig, 1968). This supply is governed by net primary productivity (NPP) and competition for resources. Geist (1987) first argued that NPP increases with latitude whilst competition decreases, permitting larger body size at higher latitudes, peaking at 60 - 65° (Wolverton et al., 2009; McNab, 2010; Huston and Wolverton, 2013). Ho et al. (2010) have also asserted that resource quality increases with latitude. |
| *Habitat availability* | Larger animals generally need larger habitats. Increased habitat fragmentation in the tropics as a result of concentrated mesoclimate gradients causes unsuitable conditions for larger species (Hawkins and Diniz-Filho, 2006; Rodríguez et al., 2008; Terribile et al., 2009). |
| *Starvation resistance* | In regions of great seasonality, larger body mass enhances fasting endurance via the allometric scaling of fat reserves (Geist, 1987; Cushman et al., 1993). |

First, Bergmann’s original hypothesis focuses on endotherms and suggests a thermoregulatory mechanism, proposing that larger organisms conserve heat more effectively in cooler environments, by virtue of their lower surface area to volume ratio, compared to smaller organisms. Therefore, a larger body size is an adaptation to cold environments because larger endotherms expend less energy per unit mass with decreasing temperature compared with smaller endotherms. Others have demonstrated that temperature seasonality can have a strong selection effect on body size patterns. Limited useful daylight time in seasonal environments can constrain species activity and impose stronger physiological limits in small bodied-species (Lindsay, 1966; Boyce, 1979).

The second mechanism relates to resource availability, which has been linked to several hypotheses. Rosenweig (1968) argued that maintenance of a particular body mass depends on a sufficient supply of food resources and to meet physiological demands, larger bodied species require more resources. He proposed that the net primary productivity of terrestrial communities increases with latitude, permitting larger sizes in higher latitudes. Giest (1987) argued that the increase in primary production with latitude permits the body sizes of herbivores to increase and consequently, the body size of their dependent carnivores. Further hypotheses have proposed that the seasonal variation of resources and in particular the amount of resources available during the reproduction and growing periods is critical for determining the body size that can be supported (McNab, 2010; Wolverton et al., 2009; Huston and Wolverton, 2011). Others, such as Lyndsay (1966) and Boyce (1979) proposed that increased seasonality of resource availability drives larger body sizes because large organisms are better able to resist longer periods of food scarcity because they have relatively more fat deposits and deplete them more slowly.

### Aims of this study

In reality, it is likely that the observed clines arise from interactions between several of these mechanisms (Mayr 1956). But one of the key challenges in resolving the causes of observed biogeographic patterns in animal body mass has been the difficulty in modelling them at regional and global scales. Although statistical modelling of empirical data is feasible, it is not an approach that can be used to resolve underlying mechanisms (i.e., correlation does not imply causation; Gaston, 2000; Marquet et al., 2014; Cabral et al., 2017). Alternative approaches that explicitly incorporate mechanisms are required. Doing so experimentally at global scales is infeasible, and hence using models based on underlying ecosystem theory and performing systematic, ‘knockout’ experiments to test the importance of individual mechanisms is a promising, yet underexplored, approach. This approach is made possible by the recent development of general ecosystem models (GEMs; Purves et al., 2013), based on fundamental ecological principles, and allowing self-assembling ecosystems to generate emergent patterns - that can be assessed against empirical data.

Here we explore the thermoregulation and resource availability mechanisms hypothesised to be responsible for latitudinal variation in endotherm body size. We apply a series of structured simulation experiments using the Madingley General Ecosystem Model (GEM), a global, virtual ecological world based on prominent ecological theories (Harfoot et al. 2014). In these simulations, macroecological patterns are entirely emergent, bottom-up, from ecological principles encoded within the Madingley model at the individual level. Therefore, we begin by evaluating the modelled biogeographic patterns in animal body mass against empirically derived patterns to demonstrate that the model can generate body size gradients that are comparable to those estimated empirically. Because of the ambiguity of latitudinal body size gradients in ectotherms, we focus on emergent endotherm patterns. Subsequently, to explore the relative importance of thermoregulation and resource availability hypotheses in driving biogeographic patterns in terrestrial animal body mass, we isolate and manipulate the model’s environmental gradients in absolute and seasonality of temperature and productivity, and examine their effects on modelled animal assemblages.

## MATERIALS AND METHODS

### The Madingley General Ecosystem Model

The Madingley model is a mechanistic and individual-based model of whole ecosystems. It was developed with the dual aims of synthesizing and advancing our understanding of ecology, and of enabling mechanistic prediction of the structure and function of whole ecosystems at various levels of organisation. It is not a statistical model built to fit data on Bergmann clines as closely as possible. Rather Madingley encodes prominent ecological theories that govern the metabolism, resource consumption, reproduction, movement and mortality and attempts to use these to generate realistic emergent properties, including latitudinal variation in community body size. It is general in the sense that it applies these same functions to all organisms in all ecosystems and individual-based because the functions are specified at the level of individual organisms. As far as we are aware it is the only model able to decompose mechanisms driving patterns in organismal body mass.

#### Pertinent components

A comprehensive description of the model is provided by Harfoot et al (2014). Here we summarise the model’s pertinent components, which simulate the dynamics of plants, and all heterotrophs with body masses above 4×10^-4^g that feed on living organisms. Organisms are not characterised by species identity and are instead grouped according to a set of categorical functional traits - for example trophic level (herbivores, omnivores and carnivores), reproductive strategy (semelparity vs. iteroparity), thermoregulatory mode (endothermy vs. ectothermy), and mobility for animals. These traits determine the types of ecological interactions that modelled organisms are involved in, whilst a set of continuous traits - total biomass of autotrophs; and current body mass, juvenile body mass, adult body mass, and optimal prey size of omnivorous and carnivorous heterotrophs - determine the rates of each process.

#### Plants

On land, plants are represented by stocks, or pools, of biomass modelled using a terrestrial carbon model. Biomass is added to the stocks though the process of primary production, the seasonality of which is calculated using remotely sensed Net Primary Productivity data (Harfoot et al., 2014). This production is allocated to above-ground/below-ground, structural/non-structural, evergreen/deciduous components. The Madingley model assumes that all above-ground, non-structural, matter is available for heterotrophic organisms to consume. Biomass is lost from plant stocks through mortality from fire and senescence, as well as through herbivory, which is described in more detail below. Production, allocation and mortality, in the plant model, are all determined by environmental conditions (temperature, number of frost days, precipitation, and the available water capacity of soils) (Smith et al., 2013).

#### Animals

Heterotrophic animals are represented in the model as ‘cohorts’, the fundamental agents of the model: collections of individual organisms occurring in the same modelled grid cell and that follow the same ecological trajectory, with identical categorical and continuous functional traits. Representing individual organisms is of course computationally unfeasible - there are simply too many individual organisms on Earth (Purves et al., 2013) - but the cohort approach enables the model to predict emergent ecosystem properties at organisational scales from individuals to the whole ecosystem. Heterotroph dynamics result from five ecological processes: metabolism, resource consumption, reproduction, mortality and dispersal (see Text S1 and Figure S1). The model can be considered as an ecological null model. It is intended to describe broad patterns, and hence does not include all ecological processes, such as, for example, ecological stoichiometry, behaviour (e.g. predator avoidance, sociality and intelligent movement) or microhabitat use. The model currently does not resolve flying organisms explicitly and so is more representative of non-volant organisms.

All endothermic functional groups in the model were iteroparous and the initial juvenile and adult body masses were drawn randomly from realistic mass ranges. The minimum endotherm juvenile mass was 0.004g, the smallest neonatal mass listed in the Pantheria database of extant and recently extinct mammal traits (Jones et al., 2009). The maximum adult mass was 5×10^6^g for herbivorous endotherms (7×10^5^g for carnivorous endotherms, 1.5×10^6^g for omnivorous endotherms) following maximal masses in these groups from the Elton Traits dataset (Wilman et al., 2014). Endothermic cohorts thermoregulate at 37°C at all times and this thermoregulation comes at no extra metabolic costs. This ecological simplification is necessary at present because in the real world the metabolic costs of thermoregulation are linked to numerous other aspects of an organism’ s ecology. Namely, behavioural responses, for which there are multiple strategies. For example, increased metabolic costs from thermoregulation in adverse environments can be avoided by hibernation or dispersal to wait for or find more clement conditions (Geiser, 2013). The model is currently behaviourally naïve in this respect and so these costs are not presently incorporated.

The model attempts to represent all organisms within ecosystems interacting with each other through dynamic and emergent trophic networks, albeit with some interactions being stronger than others. In this way, the endotherm community, and hence their median body mass, is influenced by the ecology of ectotherms in the model. Ectotherms were either iteroparous or semelparous. Cohorts were initially seeded into the model ranging between 4×10^-4^g as the smallest juvenile mass and 2×10^6^g as the largest adult mass depending on the functional group. Ectotherm activity was limited by environmental conditions following empirically-derived relationships between activity, diurnal temperature ranges, annual mean temperature and annual variation (Sunday et al., 2010). After the initial quasi-random seeding, cohorts interact with each other and with plant stocks, influenced by the environment, to select for a dynamic equilibrium ecosystem composition.

#### Simulations

The model is flexible with regard to spatial and temporal resolution. As described below we used two spatial resolutions, one at 1° x 1° to compare the model with empirical patterns, and the other at 5° x 5° for the set of knockout simulations. The coarser resolution was employed for computational efficiency given the 40 model simulations required for the knockout experiments, and as in the original formulation (Harfoot et al., 2014), we used a monthly time step throughout. We modelled the terrestrial realm exclusively, as this realm has seen the most research into empirical Bergmann clines (though see Torres-Romero et al., 2016). Comparisons to marine patterns would make an interesting further exploration. We simulated ecosystem structure for all terrestrial landmass between 65°N to 65°S in latitude. Simulations were performed on windows server machines, using this Madingley C# codebase for the simulations to generate body mass patterns for evaluations: https://github.com/mikeharfoot/C-sharp-version-of-Madingley; and, this codebase for the knockout simulations: https://github.com/mikeharfoot/Madingley-Bergmann-patterns.

### Evaluating emergent biogeographic patterns in body size

The Madingley model has previously been demonstrated to capture observed properties of individual organisms and the coarse structure of ecosystems reasonably well under environmental conditions without human impact (Harfoot et al., 2014). To explore latitudinal patterns in animal body mass, we used emergent ecosystem structure in the global grid of 1° × 1° terrestrial cells. We evaluate these against mammalian data because the model is more representative of mammals than birds, as described above. Climatological environmental conditions for each grid-cell are read as model inputs from publicly-available datasets (see Harfoot et al., 2014). For air temperature, diurnal temperature range, precipitation and number of frost days these were calculated from WorldClim mid-Holocene (approximately 6,000 years ago) downscaling of the HadGEM2-ES model reconstruction (Hijmans et al., 2005). For soil water availability, there was no equivalent Holocene climate dataset so we used average values for the period 1960 – 2000. We used the same climatological time series for each of the 100 years of model simulations, to remove the effects of inter-annual environmental variation. Importantly, we also excluded effects of anthropogenic habitat conversion and harvesting of plant or animal biomass for human use. So, our simulations represent a late-Quaternary world that has received little anthropogenic influence.

We performed 10 simulations with this protocol, in each case drawing different initial ecosystem states to capture the effects of variation in initial conditions and of stochasticity in ecological dynamics, and allowing the ecosystems to establish a quasi-steady state over a simulation length of 100 years. Figure S2 shows the temporal emergence of body mass gradients over time. It demonstrates that there is no body-mass cline associated with initial conditions. The final heterotroph biogeography is instead determined by the environmental and productivity conditions in each grid cell.

To be consistent with the empirical body size gradients (described below), which do not take abundance into account, and use species maximum or a central estimate for adult body mass, we grouped cohorts (which represent functionally equivalent, hetero-specific individuals, at a specific life stage) by their unique adult body mass, creating pseudo-species. Whilst the adult-body mass of cohorts does not change through time and so the mass of the resulting pseudo-species does not change, the composition of pseudo-species within an ecosystem can change through time. This can result from local extinction or dispersal. So, for each grid cell and each simulation we calculated the median across months of the median adult body mass of the pseudo-species present in that pixel. We calculated this using the quasi-steady state ecosystems of the final 12 months of each simulation. We took the median value across the 10 simulations as the central value across the ensemble of simulations.

### Estimating empirical biogeographic patterns of animal body mass

Estimates of the community mean body mass for mammals and birds were derived from extent of occurrence maps (EOO) and trait databases. We refer to these data as mean body mass from EOO. Extents of occurrence were intersected with the same 1° × 1° grid of cells used for the model simulations. For the species occurring in each grid cell, the median assemblage body mass was calculated (Cooper & Purvis, 2010; Rapacciuolo et al., 2017).

As introduced above, we focus our evaluation of modelled biogeographic body mass patterns on endotherms, where there is much stronger evidence of latitudinal body mass clines. For mammals, where there is evidence that anthropogenic impacts have altered the latitudinal gradients in body mass (Faurby & Araújo, 2016; Santini et al., 2017b), we used Holocene body mass data and present-natural EOO from a pre-release version 1.1 of the Phylacine database (Faurby et al 2018). The original data for body sizes mainly comes from Faurby and Svenning (2015) and Smith et al 2003, while the data on ranges mainly comes from Faurby and Svenning (2016) and IUCN (2016). These estimate what the present range would be for each mammal species given a current climate but no human impacts. The Holocene EOO maps were projected in the Behrmann equal area coordinate system so the maps were sampled at the coordinates of the 1 decimal degree model grid cell centres projected to Behrmann coordinate system. Further details can be found in Supplementary Text S2.

### Evaluating modelled biogeographic patterns of animal body mass

We evaluated emergent body mass patterns by calculating their correlation with empirical estimates across the global grid using modified t-tests to account for the loss of statistical power resulting from spatial autocorrelation (Dormann et al., 2007; Dutilleul et al., 2008). We log transformed body masses to give equal weight in the correlation to all body sizes across the orders of magnitude variation in sizes across the grid. We did not take phylogenetic correlation into account for three reasons. Firstly, because Madingley simulates cohorts of species that interact, survive or perish over time, which is different than simulating the evolution of species in phylogenies. Secondly, we focus on spatial patterns of body size rather than the time and mode in which these patterns emerge. Finally, we were not attempting to explain empirical body mass relationships with environments when phylogenetic signal should be accounted for.

### Isolating mechanisms driving body-mass patterns

#### Thermoregulation and resource availability hypotheses

We conducted a set of environmental knockout simulation experiments in the model to investigate the two major hypothesised drivers of clines in body mass: thermoregulation and resource availability. Our simulations were designed to precisely evaluate how much an environmental feature is contributing to latitudinal gradients by quantifying how much body masses change when that feature is removed from the simulations. If body mass changes considerably in response to a knockout, to the extent that the latitudinal pattern is substantially altered then we can conclude that that feature and associated mechanism plays a dominant role in driving the modelled body size patterns. We considered two aspects of the thermoregulatory hypothesis. First, that the latitudinal gradients in annual mean temperature (mean monthly temperature across the year) selects for larger endotherms where annual mean temperature is lower. Second that variation in temperature through the year (seasonality of temperature) selects for larger body mass where seasonality of temperature is higher. We also considered whether these conditions select for smaller ectotherms, through their capacity to more effectively utilise benign windows in otherwise unsuitable environments (e.g. Olalla-Tárraga and Rodríguez, 2007). Analogously to the temperature knockout simulations, we tested for the effects of latitudinal gradients in the cumulative annual net primary production and the variation in net primary productivity across months of the year (seasonality of production). Our prediction was that higher annual mean productivity (mean monthly productivity across the year) and greater seasonality of production selects for larger endotherms (Lindsey 1966, Rosenweig, 1968, Boyce 1978, 1979). We therefore conducted, in addition to a full (control) simulation with no environmental feature changed, four knockout experiments in each of which we removed either a gradient or variation in one of these aspect of the environment at a time (Table 2, which lists treatment names, Fig S3). These knockout simulations were then compared to the full simulations.

**Table 2.**
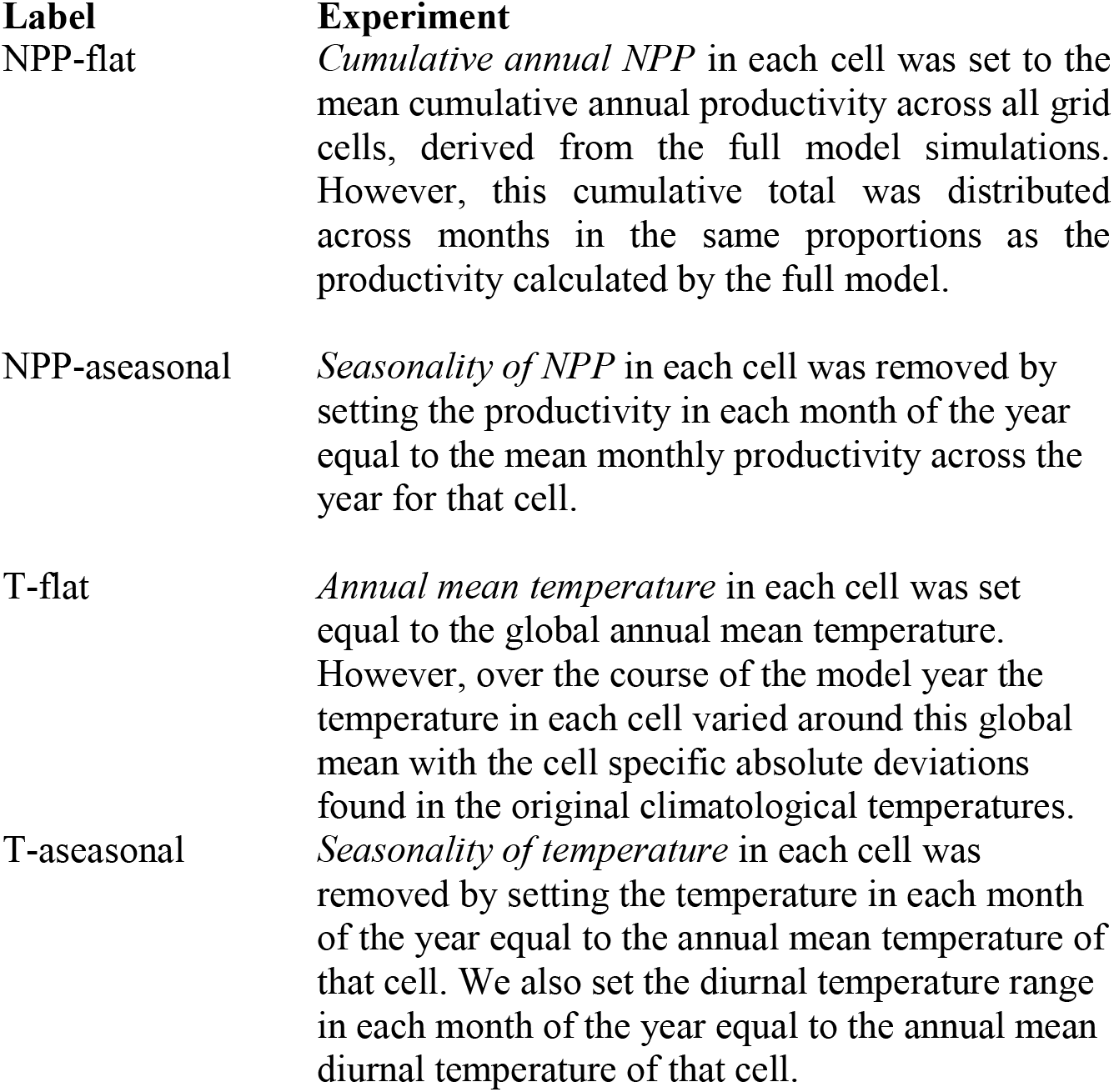
Environmental knockout experiments

| **Label** | **Experiment** |
| --- | --- |
| NPP-flat | *Cumulative annual NPP* in each cell was set to the mean cumulative annual productivity across all grid cells, derived from the full model simulations. However, this cumulative total was distributed across months in the same proportions as the productivity calculated by the full model. |
| NPP-aseasonal | *Seasonality of NPP* in each cell was removed by setting the productivity in each month of the year equal to the mean monthly productivity across the year for that cell. |
| T-flat | *Annual mean temperature* in each cell was set equal to the global annual mean temperature. However, over the course of the model year the temperature in each cell varied around this global mean with the cell specific absolute deviations found in the original climatological temperatures. |
| T-aseasonal | *Seasonality of temperature* in each cell was removed by setting the temperature in each month of the year equal to the annual mean temperature of that cell. We also set the diurnal temperature range in each month of the year equal to the annual mean diurnal temperature of that cell. |

#### Experimental protocol

To account for stochastic effects, we performed an ensemble of 10 simulations for the control conditions and for each knockout, which yielded 50 simulation runs. It was computationally unfeasible to run these simulations at a resolution of 1° × 1°, so we used a coarser resolution of 5° × 5° cells with the same extent. The change in resolution resulted in consistent variation body mass across latitude, however in general the annual median, community median body size was larger in the coarser resolution simulations (Fig S4). For each knockout experiment, we used the median across the ensemble as the central estimate. For each knockout across each latitude band, we then calculated the relative median body mass deviation from the unmodified control simulation. Because the response of ectotherms can impact on that of endotherms, we include their response to environmental knockouts in the studies.

#### Exploring causation

To further explore the causes of changes in latitudinal body mass clines in response to environmental knockouts, we analysed changes in individual level feeding, reproduction and mortality rates. We ran simulations for two grids of four 5° × 5° cells, one in North Asia (55-65°N and 75-85°E) and one in equatorial Africa (5°S-5°N and 5-15°E). These grids were representative of different environments and exhibited analogous patterns to the global simulation when run outside the global model. These simulations were run for 20 years and for each month we exported the process rates (feeding/assimilation, reproductive output, per capita mortality) of every cohort as well as cohort properties (e.g. current body mass, adult mass and cohort abundance). It was not possible to run a global simulation because of the prohibitively long runtimes and large volumes of individual level ecological data generated. Annual rates of the ecological processes were calculated for each individual within each cohort by summing the monthly outputs. See Text S3 for further methodological details.

## RESULTS

### Global patterns of modelled animal body mass

Spatial patterns of endotherm and ectotherm body mass emerge from the model without exogenous constraints based on its parsimonious descriptions of the ecological processes governing ecosystem structure and function (Figure S2). These patterns exhibit strong spatial and latitudinal variation (Figure 1). For endotherms, median body mass generally increases with aridity and with latitude (Figure 1a).

**Figure 1.**
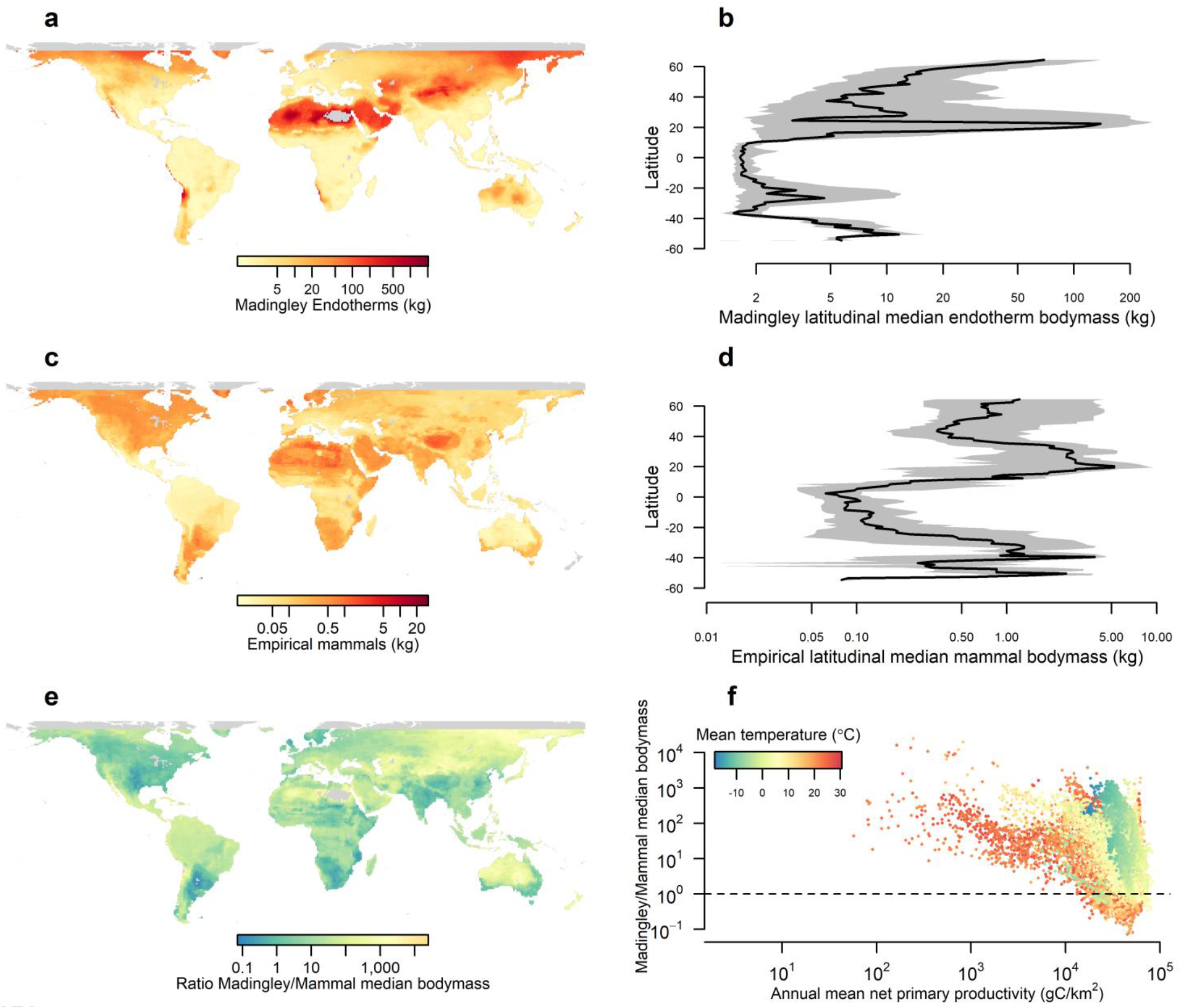
Global distributions (a, c) and latitudinal profiles (b, d) of ensemble median, annual mean, community median body masses of endotherms predicted by the Madingley model (a, b) and Holocene mammals estimated from empirical data (c, d). The ratio of Madingley to empirical estimates is plotted spatially (e) and the pixel values plotted as a function of the annual mean net primary productivity and coloured according to the annual mean temperature (f). Black line in b and d indicate the median body mass across each latitude band, whilst the grey shading indicates the interquartile range.

### Comparing modelled and empirical body mass

#### Endotherms

Endotherms modelled by Madingley exhibit spatial variation in body mass that resembles empirical patterns (Figure 1a, c). Modelled body sizes increased in northern USA and Canada, south west Africa, the Sahara, and central Asia consistent with empirical patterns in mammals. Modelled and empirical body sizes were significantly, positively correlated at the global scale (Mammals: r = 0.38, p < 1 ×10^-5^, modified t-test). The ratio of modelled and empirical body mass estimates shows many areas in the world with consistent absolute body mass values (Figure 1e). However, northeast Asia, central and western Australia stand out as showing disagreement. In these areas, the Madingley model suggests larger body masses than the east coast of Australia, whilst the empirical data for mammals suggest the opposite.

Latitudinal profiles show that Madingley predicts Bergmann-like clines that are similar in shape to those for Holocene mammals (Figure 1b, d). Body masses reached a minimum in the sub-tropics. They increased towards the highest latitudes but with peaks around the desert belt. Although modelled and empirical body masses are correlated, in general, the organisms predicted by Madingley were larger than those estimated from empirical data by about an order of magnitude.

There was a trend in the degree of agreement between modelled and empirical body mass estimates as a function of NPP (Figure 1f). As annual mean NPP decreased, Madingley tended to increasingly overestimate endotherm body mass. This relationship held across different temperature regimes. Agreement was more likely in locations of higher NPP but there was nonetheless substantial variation in the ratio of modelled to empirically estimated body mass for high NPP locations even within similar annual mean temperature environments.

### Knockout experiments

#### Environmental knockouts

We found support for the resource availability drivers of body mass clines and in particular the seasonality of resource availability (Figure 2). When the latitudinal gradient in NPP was removed (NPP-flat, Table 2), endotherm body masses modestly but significantly decrease (negative proportional body mass deviation, with 95% interval below zero) across most latitudes, so there was little change in the body-mass clines (Figure S5). However, when we removed variation in NPP across the year (NPP-aseasonal), body masses declined significantly in high latitudes of the northern hemisphere, with an effect size that increased with increasing latitude. Changes were of smaller magnitude and less frequently significant in tropical latitudes with the result that body mass tended to decline with increasing latitude in the northern hemisphere, almost removing the conventional Bergmann clines (Figure S5). In both NPP-flat and NPP-aseasonal knockouts, the latitudinal median of the grid-cell median adult endotherm body mass was reduced (median declines of 21% for NPP-flat and 27% for NPP-aseasonal; Figure 2). Removing variation in temperature across the year (T-aseasonal) caused body size declines in mid-latitudes in both hemispheres and had little effect in low latitudes. Effect sizes were smaller and less significant in the southern hemisphere for all knockouts. T-flat had little effect on endotherm body mass in the either hemisphere.

**Figure 2.**
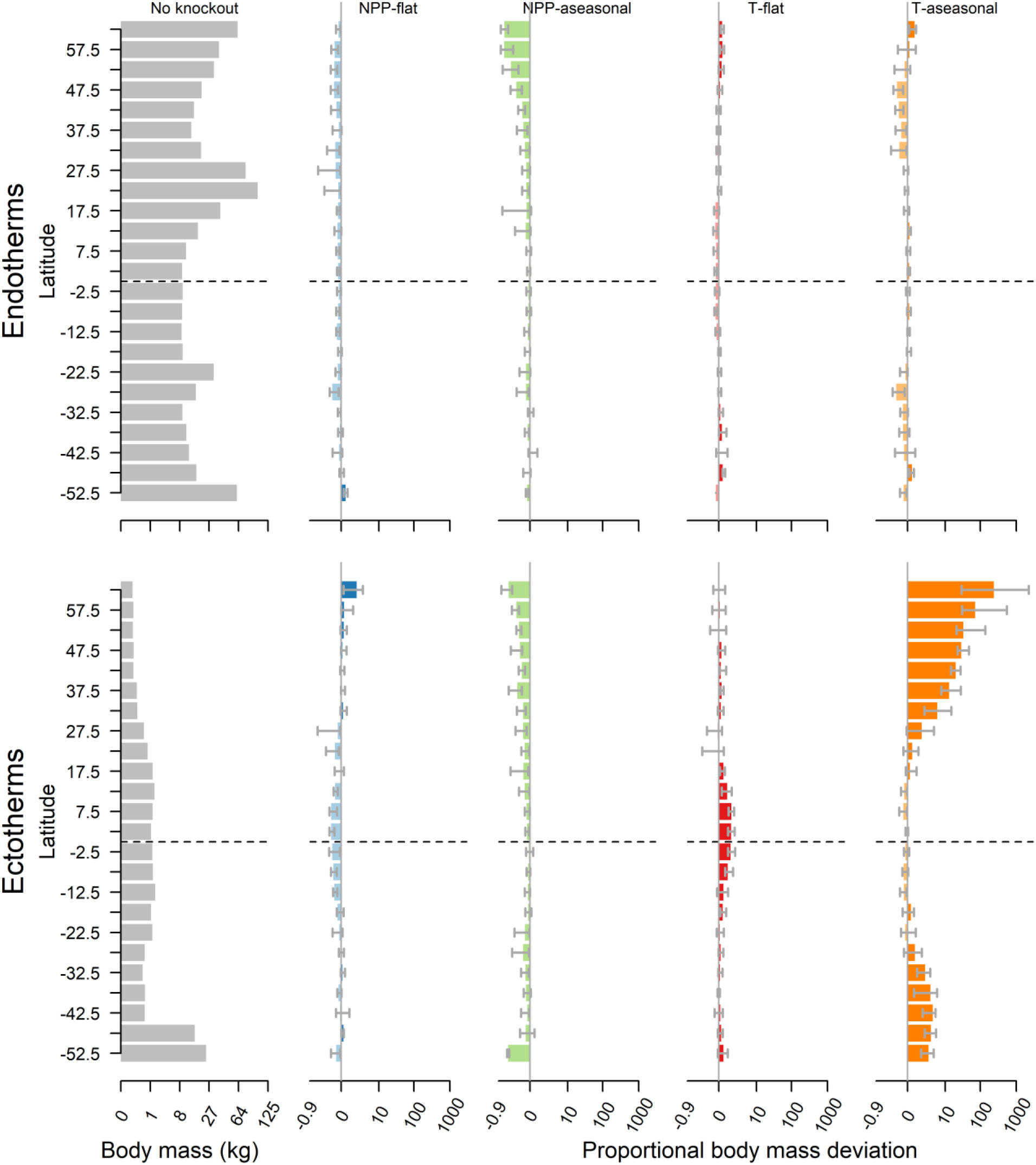
The relative effect on community mean endotherm and ectotherm body mass across latitudes of removing spatial variation in cumulative annual primary production, seasonality of primary production, annual mean temperature and annual temperature seasonality. Solid bars represent the median and grey lines the interquartile range across longitudes with each latitude band. Lighter hue indicates negative median effect whilst darker indicates a positive median effect

Ectotherms body mass tended to be greater in the tropics and decline with increasing northern latitude. In response to the knockouts, this body mass cline responded most to seasonality of resource availability and of thermoregulation. Ectotherm body masses clines were exacerbated when there was no seasonal variation in NPP (NPP-aseasonal) but reversed when there was no seasonal variation in temperature (T-aseasonal) (Figure 2 and Figure S5).

#### Exploring causation: Individual level ecological responses

The results from a grid cell in North Asia (central Russia, 62.5°N, 82.5°E) showed that, for endotherms, in all knockouts other than T-flat, median body masses declined because smaller organisms persisted in the model (Figure 3a). NPP-aseasonal had the largest effect followed by T-aseasonal. In terms of differential effects on organisms of different body mass, smaller organisms tended to assimilate more biomass, produce more offspring and have lower starvation mortality losses relative to the full environment for all knockouts except T-flat (Figure 3b, S5).

**Figure 3.**
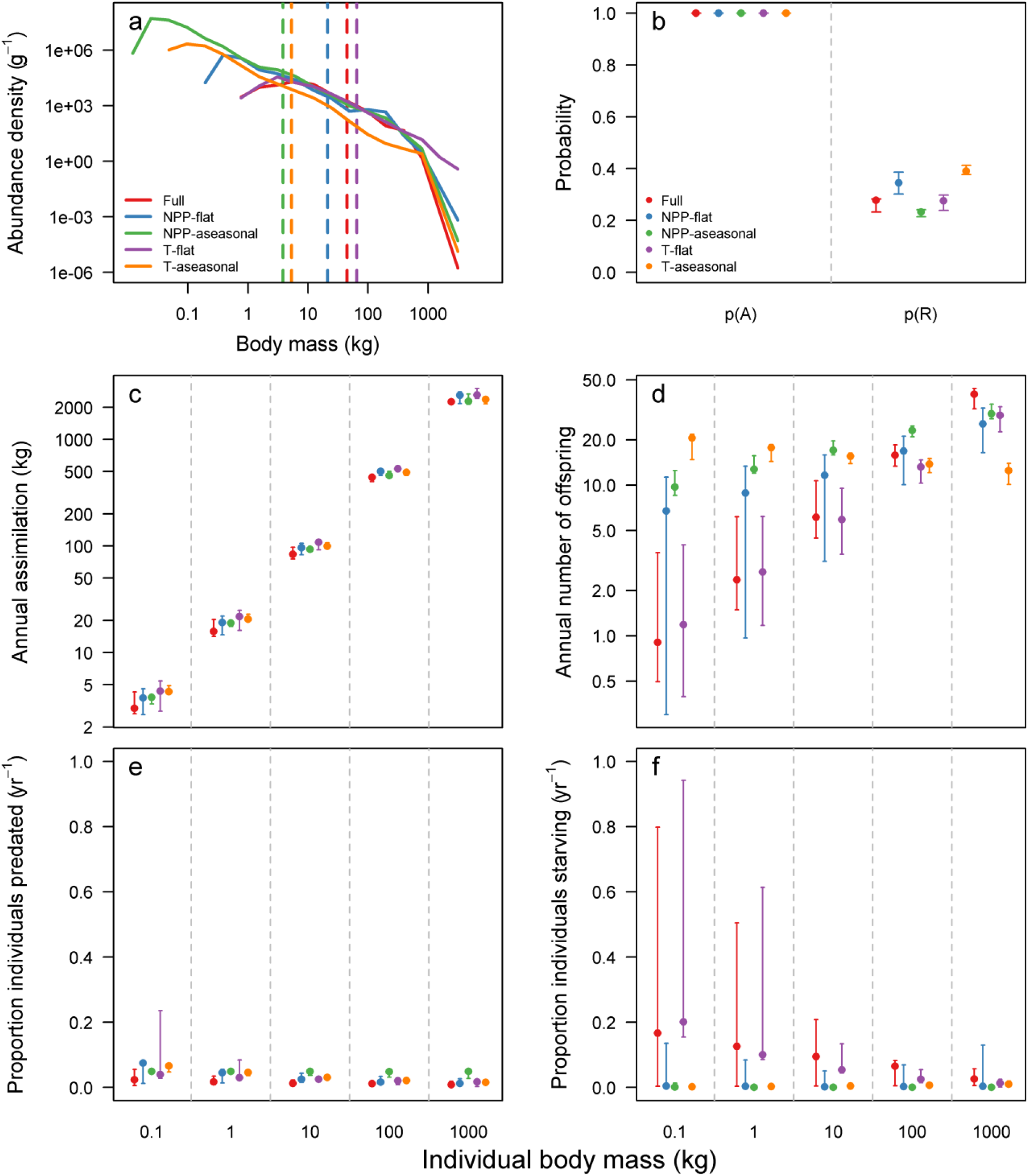
Effect of environmental knockouts in a grid cell centred at 62.5°N and 82.5°E on (a) endotherm community size distribution, (b) probability of assimilating biomass, p(A), and of reproducing offspring, p(R), within a year, (c) annual mass assimilated, (d) number of offspring produced per individual present at the start of the year, and proportion of individuals present at the start of the year that die from predation (e) or starvation (f) events. The community size distribution is the annual median abundance per mass bin from the 20^th^ year of the simulation, dashed lines indicate the community mean body mass for each knockout. Points in b – f represent median predicted values from models fitted to annual data from the last 5 years of a 20-year simulation, error bars represent maximum and minimum value across years.

For ectotherms, median body mass increased in all knockout simulations except the NPP-flat (Figure 4). The largest effects arose in the T-flat and T-aseasonal simulations. T-flat resulted in ectotherms with body mass less than 10g dying out, and orders of magnitude lower abundances of larger ectotherms. This arose because the interaction of warmer temperatures but with retained seasonality patterns reduced the period of the year in which ectotherms were active in the model. As a result, the likelihoods and rates of assimilation and reproduction were reduced and rates of starvation mortality were substantially elevated, especially for smaller ectotherms with a higher relative metabolic rate. T-aseasonal resulted in decreased abundance of the smallest, and an increase in the abundance of the largest, ectotherms. Ectotherms in this experiment were active all year round in the model (there was no unsuitable winter season) hence the probabilities and rates of assimilating or reproducing raised and reproductive output was increased, especially for smaller organisms.

The results from environmental knockouts in an equatorial ecosystem (2.5°N and 12.5°E) showed limited changes in the abundance size distribution of endotherms (Figures S6-S8). Conversely, for ectotherms, the median body mass decreased by five orders of magnitude in the T-aseasonal knockout. This was caused by the increased likelihood and rate of reproduction of smaller ectotherms but reduced output of larger ectotherms, compared to the full environment. Smaller ectotherms were also less likely to starve in the aseasonal temperature environment when compared to the full environment.

**Figure 4.**
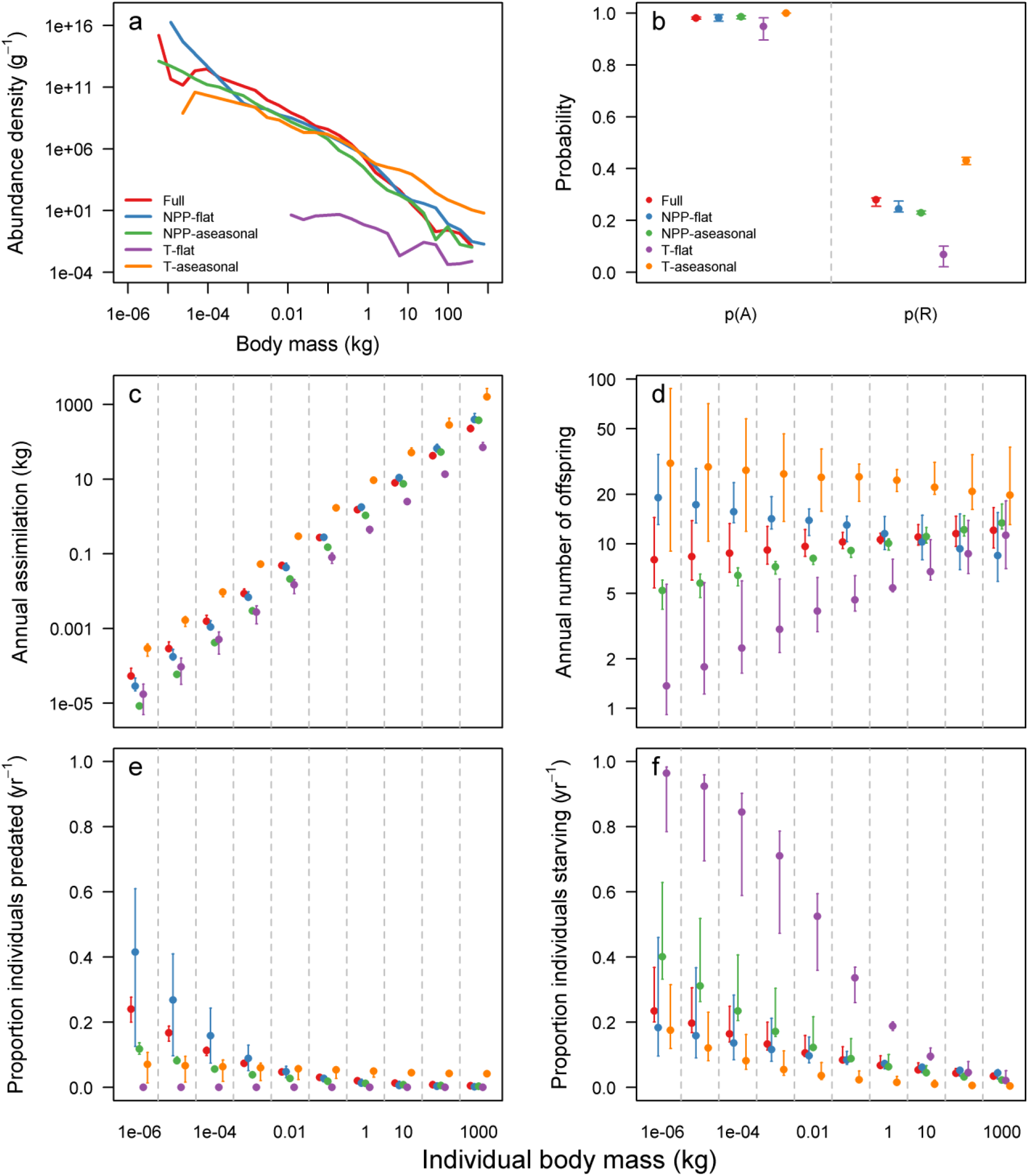
Effect of environmental knockouts in a grid cell centred at 62.5°N and 82.5°E on (a) ectotherm community size distribution, (b) probability of assimilating biomass, p(A), and of reproducing offspring, p(R), within a year, (c) annual mass assimilated, (d) number of offspring produced per individual present at the start of the year, and proportion of individuals present at the start of the year that die from predation (e) or starvation (f) events. The community size distribution is the annual median abundance per mass bin from the 20^th^ year of the simulation, dashed lines indicate the community mean body mass for each knockout. Points in b – f represent median predicted values from models fitted to annual data from the last 5 years of a 20 year simulation, error bars represent maximum and minimum value across years.

## DISCUSSION

Bergmann’s rule is an ecological pattern for which there is a complex set of hypothesised mechanisms and varying levels of empirical support. The Madingley model is a general mechanistic model of whole ecosystems, it was not built to fit Bergmann cline data specifically, rather to model emergent patterns based on encoded ecology. Encouragingly, the model is able to generate global spatial and latitudinal patterns in body mass from individual organismal interactions that are consistent with empirical estimates. Attempting to unpick the variation in body-size attributed to different environmental and ecological drivers provides insight into the potential mechanisms underlying empirical observations. In general, the model captures endothermic patterns reasonably well. Madingley generally over-predicted the body size of organisms compared to empirical estimates. There are several possible factors that might contribute to this disagreement.

First, the median unique adult body masses for all cohorts in a grid cell is not equivalent to the median species mass. Because the cohorts in Madingley are hetero-specific, at present the model does not inform about how many species it might represent. A cohort of small endotherms in Madingley might represent many species with small body size. Whilst several cohorts of large organisms in Madingley, might represent intra-specific variation in a single species. Second, the empirical estimates have substantial associated uncertainties. For example, there can be considerable intraspecific variation in body size across ranges (Ashton, 2002; Clauss et al., 2013; Tseng and Soleimani Pari, 2019). The EOO maps assumes a species is found in every cell within its EOO at all times. Thirdly, the Madingley model might be missing mechanisms that limit the maximum size of organisms or increase the survival of smaller organisms. The current formulation of the model misses several aspects of thermoregulatory behaviour. For example, ectotherms can behaviourally regulate their temperature above or below that of the ambient temperature (Kearney et al., 2009), a process not included in Madingley. So metabolic costs may be under-estimated in cold environments and over-estimated in hot environments. In addition, the model neglects hibernation, and so metabolic costs might be over-estimated in seasonal environments. Including such behaviours could alter the degree of converse Bergmann cline in the model but it is not clear in which direction. The model also currently assumes that thermoregulation in endotherms has a flat metabolic cost across latitudes and that there is no extra metabolic cost associated with, for example, fat deposits, feathers or fur to protect against cold temperatures. Since smaller endotherms pay a greater relative metabolic cost to thermoregulate against extreme temperatures than larger endotherms (Porter and Kearney, 2009), including this effect would likely amplify the latitudinal body mass gradients in the model. As a result of their possibly opposing effects and the complex community assembly processes operating through time, the net impact of including thermoregulatory effects on community body masses is not clear, but should be explored as these features are incorporated into the model. The tendency for the Madingley model to overpredict endotherm body size in lower productivity environments (Figure 1 e, f) provides evidence for missing mechanisms, such as hibernation, that permit smaller organisms to survive adverse conditions, or that prevent larger organisms outcompeting smaller ones, for example as a result of resources being inaccessible to larger organisms. The lack of three dimensional structure to the vegetation in the model is one example of the latter, arboreal mammals tend to be smaller bodied than ground-based mammals. An additional omission that might impact the largest body sizes in the model is the absence of pack hunting, which would enable predators to predate on larger prey, whilst currently predators are generally larger than their prey organisms.

More generally, the patterns that emerge from the model will be sensitive to uncertainties in the model structure (which ecological processes are represented and how they are formulated) and the parameterisation of these functional forms. Exploring this sensitivity should be a critical next step, as this would permit an exploration of the generality of macroecological mechanisms responsible for Bergmann clines. Notwithstanding this caveat, the results from the current, published and evaluated version of the model results provide support for the role of both thermoregulation and resource availability mechanisms, and particularly the seasonality of these, as key mechanisms determining body mass of both endotherms and ectotherms.

Despite being hugely simplified in comparison to reality and attempting to model all ecosystems on land at the level of individual organisms, the fundamental ecological mechanisms currently included in the Madingley model produce plausibly realistic emergent geographic pattern of median endotherm body mass. It also captures much of the variation of median body mass within each latitudinal band. For example, desert regions are known as prominent sites of large body mass for birds and mammals (Blackburn and Hawkins, 2004; Olson et al., 2009; Morales-Castilla et al. 2012a), and this is produced by the model outputs. The Saharan and Gobi deserts all harbour greater median animal body mass than the median of their respective latitudes (Figure 1).

We tested hypotheses on the mechanistic factors behind observed patterns through knock-out experiments for environmental factors. The initial conditions of the model are spatially neutral, meaning that there is no initial spatial pattern in the types of cohorts seeded in each grid cell. Therefore, the model does not require chance or mechanisms of migration ability or historical contingencies to generate body mass gradients (see e.g. Morales-Castilla et al. 2012b), implying that while these hypothesised mechanisms may be responsible, Bergmann’s clines can be produced without them. There is no *a priori* latitudinal variation in predation risk in the model, therefore this mechanism is also not required for the emergent body mass patterns (Wolverton et al., 2009). Our results suggest that thermoregulation and resource availability are dominant direct controls on endotherm community mean body mass, they as when these factors are removed, or “knocked out”, from the model, endotherm body mass across latitudes declines more than for other factors. Where productivity increased or the seasonality of productivity was removed, the severity of the seasonal cycling of food surplus and deficit (sensu Geist, 1987) was reduced, allowing smaller organisms that are more susceptible to starvation, to avoid this risk and increase their fecundity. The resulting abundance increase of smaller endotherms thus draws the community mean body mass down and also exerts some bottom up control over larger endotherms, which have higher starvation rates and lower reproduction rates than the full environment simulations.

For modelled ectotherms, thermoregulation is the dominant control on community body mass through its effect on enabling access to resources. This is because ectotherm activity is limited under extreme ambient temperatures. In simulations that isolated the seasonality of productivity from that of temperature (NPP-aseasonal vs T-aseasonal), the altered seasonal cycling of food surplus and deficit appears unimportant if the temperature remains seasonal such that the organism’s ability to use those resources is impeded.

General Ecosystem Models, such as the Madingley model, can generate ‘in-silico’ global ecosystems that can be experimentally manipulated in ways that are impossible in the real world - allowing for novel explorations in ‘mechanistic macroecology’. Our study demonstrates this for Bergmann’s rule, and provides support for the thermoregulation and resource availability hypotheses, while also generating an entirely new hypothesis of trophic interactions between ectotherms and endotherms as a potential structuring mechanism for observed clines in endotherm body mass. While ‘mechanistic macroecology’ cannot be the only solution or approach to unpicking global macroecological patterns, our study suggests that it is a tool that should be investigated further beyond these preliminary explorations, and can provide both ecological insight and identify model weaknesses or missing processes. Furthermore, our finding of a bottom up control of endotherm body mass by ectotherms suggest that we must model ecosystems and biological interactions holistically to better capture how they are structured and function.

## DATA ACCESSIBILITY

All modelled body mass data has been deposited in Data Dryad (DOI to be added on acceptance). Empirical data are accessible from the cited sources.

