## Supplementary figures and tables for "Mechanistic macroecology: exploring the drivers of latitudinal variation in terrestrial body size in a General Ecosystem Model" (from another version of this preprint: version 1, 2019-09-19)

**Text S1. Details of ecological mechanisms**

Predator-prey interactions (including herbivory) are based on a Holling's Type III functional response (Denno et al., 2012), and for predation on a size-based model of predator-prey feeding preferences (Williams et al, 2010). Metabolism is based on empirical relationships between energy consumption and ambient temperature taking into account the body mass of the organism (Brown et al., 2004). Endotherms are assumed in the model to thermoregulate perfectly, and thus are active for 100% of each time step. Ectotherms in the model do not thermoregulate, and thus are only active for the proportion of each time step during which ambient temperature was within their upper and lower activity temperature limits, estimated following Deutsch et al. (2008). Reproduction can occur once a cohort has achieved its adult body mass and results from the allocation of surplus mass to reproductive potential followed by reproductive events once a threshold ratio of reproductive potential to adult body mass is reached (Harfoot et al., 2014). Mortality (in addition to predation mortality) arises from three causes: a constant background rate, starvation if insufficient food is obtained, and senescence, which increases exponentially after maturity with a functional form similar to the Gompertz model (e.g. Pletcher, 1999). Dispersal in the terrestrial realm is either random diffusive dispersal of juvenile organisms or directed dispersal of organisms in response to starvation (i.e. dispersal away from unsuitable feeding conditions) or low densities of individuals (i.e. dispersal to find a mate when encounter rates are very low) (Harfoot et al, 2014).

**Text S2: Methodological details for calculation of mean body masses from extent of occurrence maps**

*Excluding marine mammals*

Marine mammals were excluded from the extent of occurrence and trait datasets by excluding all species from the following families: Otariidae, Odobenidae, Phocidae, Trichechidae, Dugongidae, Physeteridae, Kogiidae, Monodontidae, Ziphiidae, Delphinidae, Phocoenidae, Platanistidae, Iniidae, Pontoporiidae, Balaenidae, Neobalaenidae, Balaenopteridae and Eschrichtiidae.

*Community median ectotherm body masses*

For reptiles and amphibians, historic range maps are not available. So present day EOO maps were taken from the IUCN Red List (IUCN, 2017) for amphibians and from Roll et al. (2017) for reptiles. For reptiles we used maximum body masses available for 9,631 species estimated using body lengths and allometric scaling relationships (Meiri, 2010; Feldman and Meiri, 2013; Feldman et al., 2016). Body mass data for 5,748 amphibians were extracted from AmphiBIO (Oliveira et al. 2017) using empirical masses (=2011) or estimated (=3737) using body length and allometric relationships (Santini et al., 2017a).

*Estimating amphibian body masses using allometric scaling relationships*

We used data from Oliveira et al., 2017 and Santini et al., 2017 on snout to vent lengths and body mass to derive allometric relationships between SVL and body mass for anurans, caudatans, caecilians (order gymnophiona). We fitted separate models for each order following the evidence in Santini et al., 2017 supporting fitting the orders separately. We did not include information on the habitat preferences for species since this was not available for most species for which SVL and body mass were available. Both SVL and body mass were $\log_{10}$ transformed prior to linear model fitting. Model statistics show that the models capture the data reasonably well ($r^2 > 0.6$; $p < 0.01$), despite the fact that there was limited data available for caecilians (Table S1).

**Text S3: Methodological details for analysis of individual level ecological process responses to environmental knockouts.**

We were interested in how the relationship between annual rates of the ecological processes (feeding/assimilation, reproductive output, per capita mortality) and the body mass of the individuals varied with respect to environmental knockout simulations. Since the body mass of individual cohorts is a dynamic variable in the model, whilst their mass at maturity is fixed, we used the latter to describe the mass of the cohort. We used log-transformed adult mass to fit a linear relationship with each process rate. For reproduction and feeding some individuals had zero rates (i.e. starved and were reproductively unsuccessful), so we used a hurdle modelling approach: We first calculated the probability of a non-zero rate using logistic regression for each rate. Subsequently, for individuals with non-zero feeding or reproductive rates, we then performed a gamma regression with an identity link function to estimate the relationship between the process rate and the log transformed adult mass. For both predation and non-predation mortality, we used generalised linear models to describe the relationship between the dependent variable mortality rate and independent variable the log-transformed adult mass. Since the mortality rates were overdispersed we used a quasibinomial link function.

Table S1 Summaries for models for allometric scaling between $log_{10}$ transformed snout to vent length and $log_{10}$ transformed bodymass for the amphibian orders, anura, caudata and gymophiona.

| Group | Intercept estimate | Slope estimate | P value | Adjusted $r^2$ | Sample size, N |
|---|---|---|---|---|---|
| Anura | -3.29 | 2.47 | < 1E-5 | 0.69 | 580 |
| Caudata | -3.07 | 1.83 | < 1E-5 | 0.62 | 113 |
| Gymnophiona | -13.69 | 5.75 | 0.003 | 0.66 | 10 |

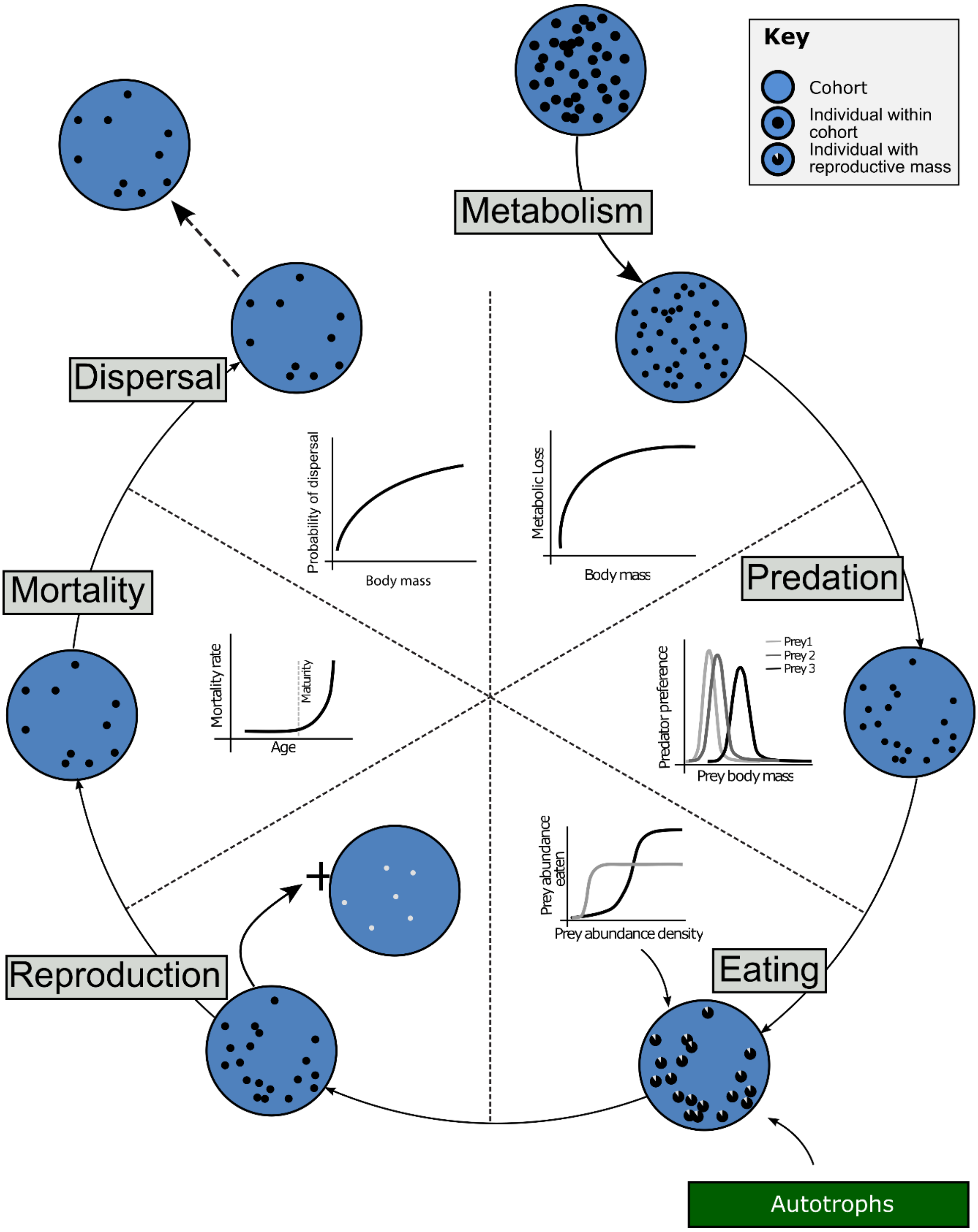


**Figure S1:** Modelled ecosystem structure and function emerges from a combination of processes operating on individual organisms within a grid cell.

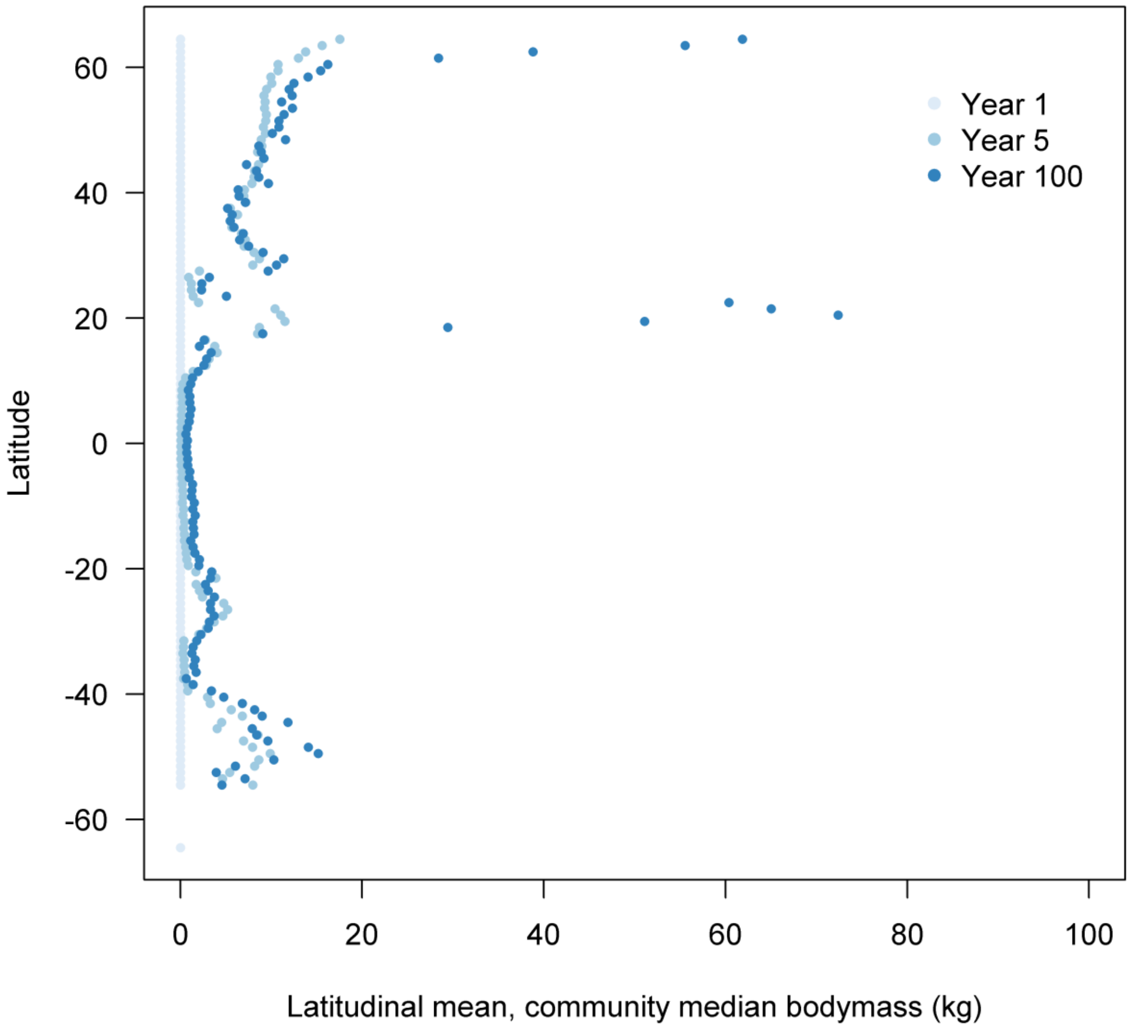


**Figure S2:** Emergence of endotherm latitudinal body mass patterns over time from neutral initial conditions.

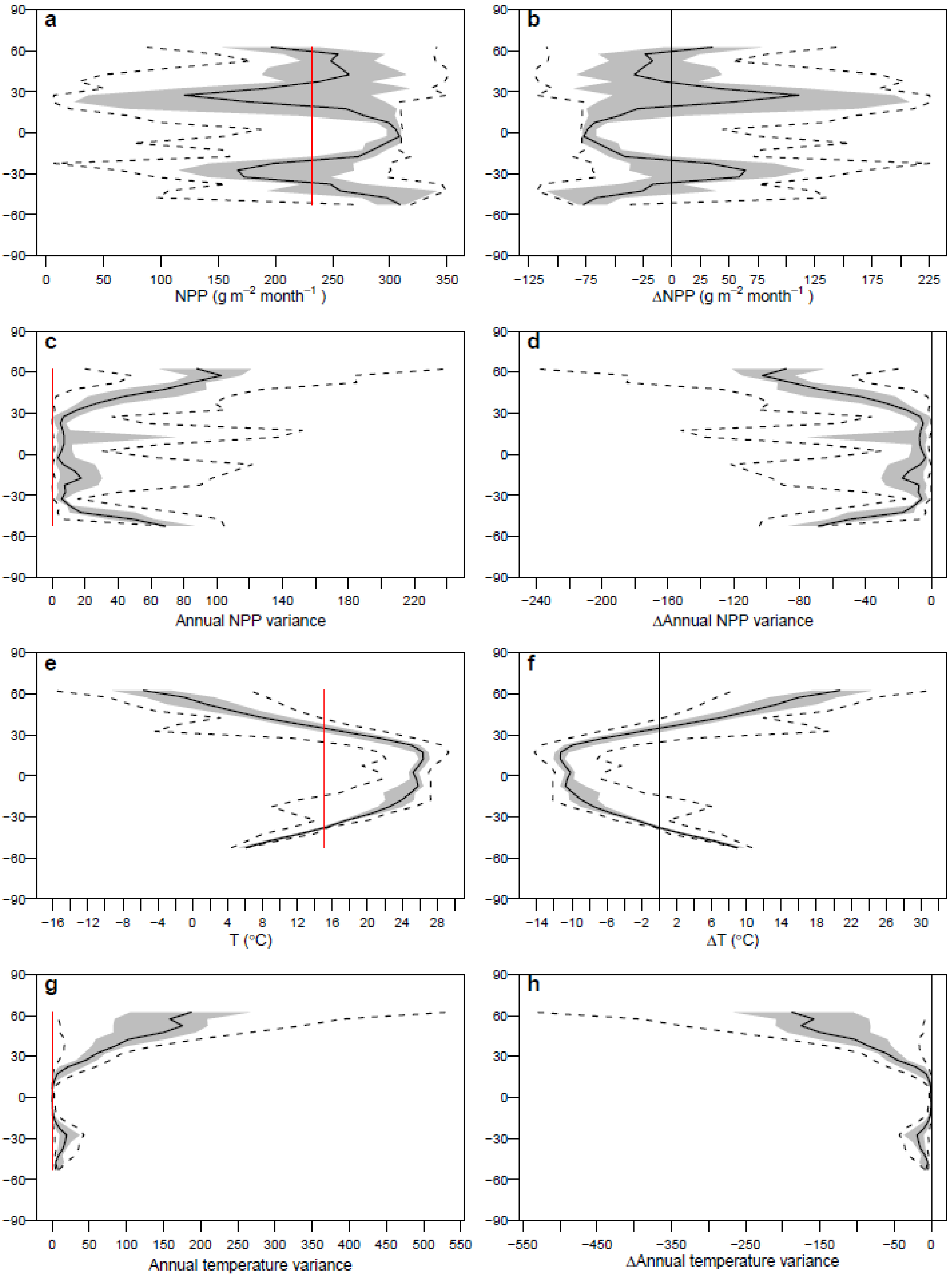


**Figure S3.** Latitudinal profiles of environmental conditions for the full model and for the knockout experiments: annual mean net primary productivity for the full model (a) and the change in annual mean net primary productivity under NPP-flat conditions (b); annual NPP variation for the full model (c) and the change in annual NPP variation under NPP-aseasonal conditions (d); annual mean temperature for the full model (e) and the change in annual mean temperature under T-flat conditions (f); annual mean temperature variation in the full model (g) and the change in annual mean temperature variation under T-aseasonal conditions (h). Solid profile like reflects the median value across longitudes in each latitude band, the grey shading represents the interquartile range and the dashed lines reflect maximum and minimum values. Vertical red lines in a, c, e, g represent the value for the respective knockout conditions. Vertical black lines in b, d, g, h mark zero on the x-axis.

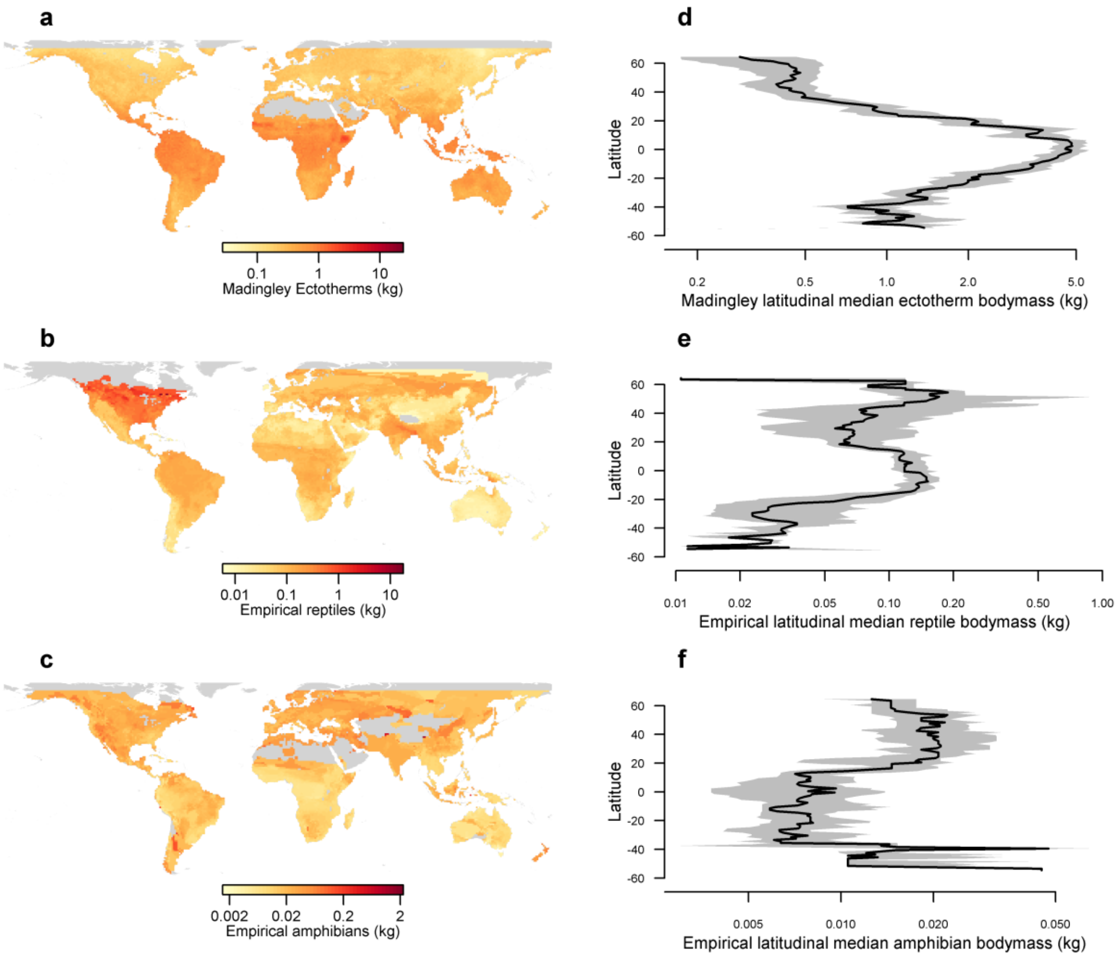


**Figure S4.** Global distributions (a-c) and latitudinal profiles (d-f) of annual mean, community median body masses of ectotherms predicted by the Madingley model (a, d), present-day reptiles estimated from empirical data (b, e), present-day amphibians estimated from empirical data (c, f).

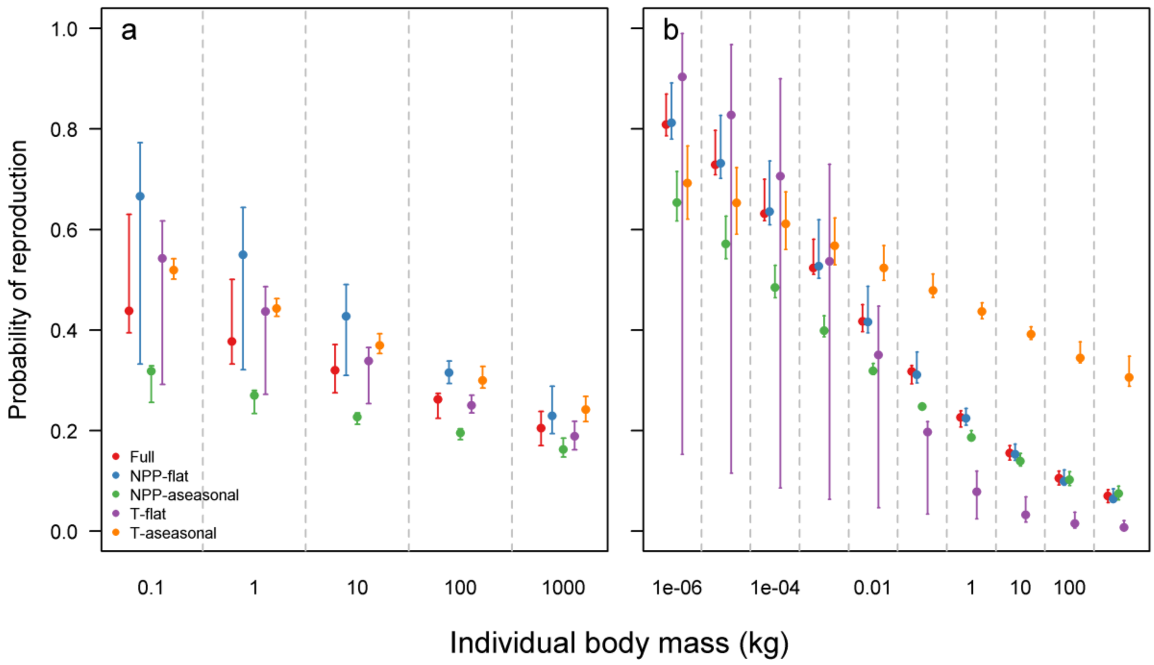


**Figure S5.** Probability of reproduction as function of body mass for endotherms (a) and ectotherms (b) for a grid cell centred at 62.5°N and 82.5°E

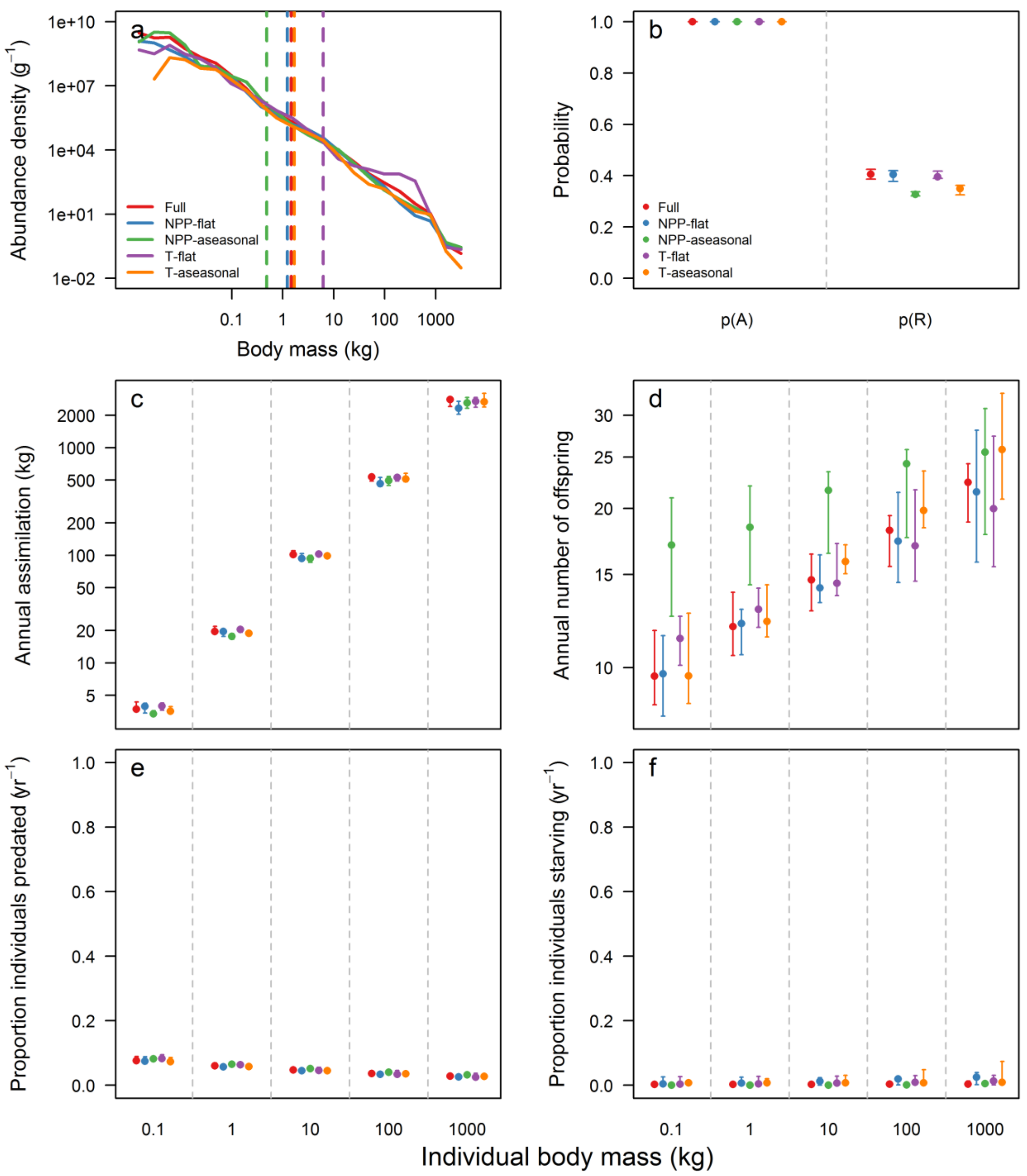


**Figure S6.** Effect of environmental knockouts in a grid cell centred at 2.5°N and 12.5°E on (a) endotherm community size distribution, (b) probability of assimilating biomass, p(A), and of reproducing offspring, p(R), within a year, (c) annual mass assimilated, (d) number of offspring produced per individual present at the start of the year, and proportion of individuals present at the start of the year that die from predation (e) or starvation (f) events. The community size distribution is the annual median abundance per mass bin from the 20$^{th}$ year of the simulation, dashed lines indicate the community mean body mass for each knockout. Points in b – f represent median predicted values from models fitted to annual data from the last 5 years of a 20 year simulation, error bars represent maximum and minimum value across years.

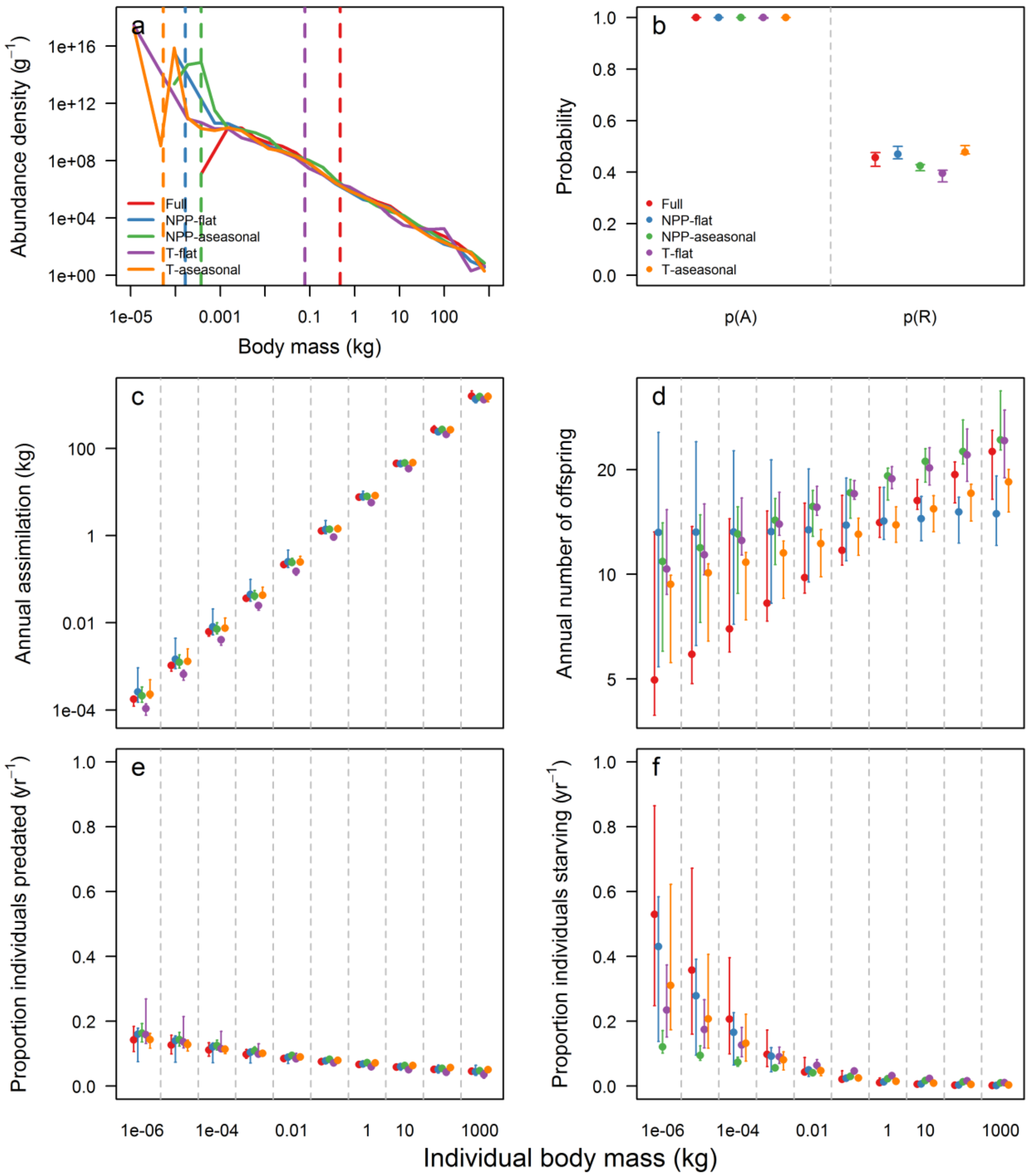


**Figure S7.** Effect of environmental knockouts in a grid cell centred at 2.5°N and 12.5°E on (a) ectotherm community size distribution, (b) probability of assimilating biomass, p(A), and of reproducing offspring, p(R), within a year, (c) annual mass assimilated, (d) number of offspring produced per individual present at the start of the year, and proportion of individuals present at the start of the year that die from predation (e) or starvation (f) events. The community size distribution is the annual median abundance per mass bin from the 20th year of the simulation, dashed lines indicate the community mean body mass for each knockout. Points in b – f represent median predicted values from models fitted to annual data from the last 5 years of a 20 year simulation, error bars represent maximum and minimum value across years.

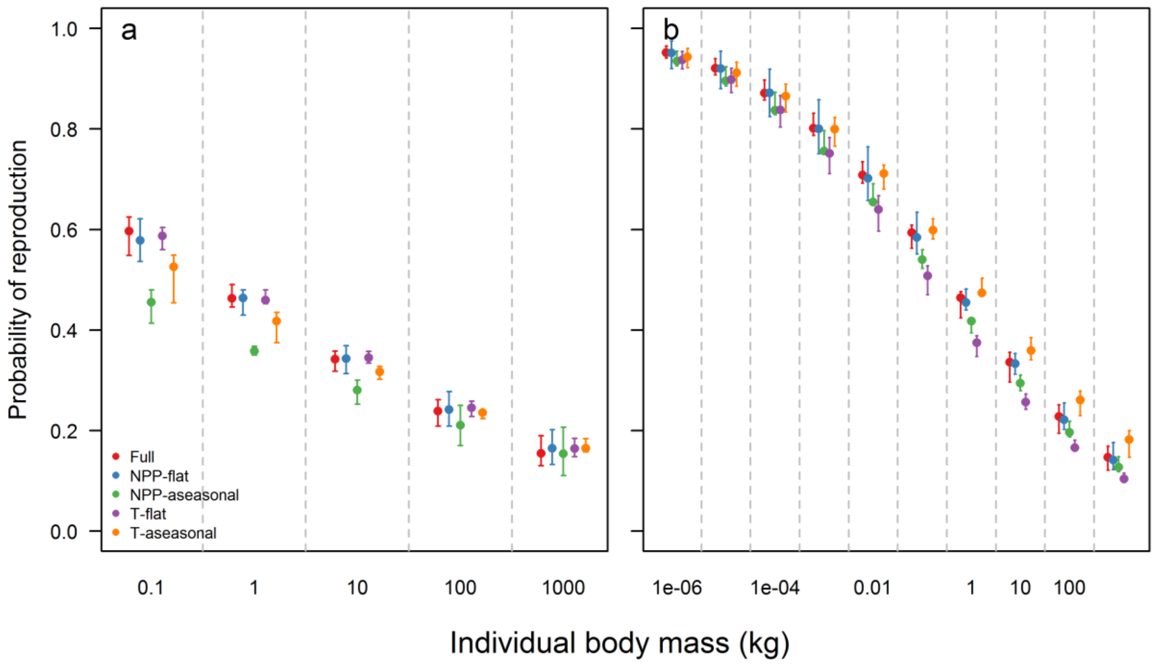


**Figure S8.** Probability of reproduction as function of body mass for endotherms (a) and ectotherms (b) for a grid cell centred at 2.5°N and 12.5°E
